# Combination drug screen targeting glioblastoma core vulnerabilities reveals pharmacological synergisms

**DOI:** 10.1101/2022.12.14.520491

**Authors:** Jérémy Ariey-Bonnet, Raphael Berges, Marie-Pierre Montero, Baptiste Mouysset, Patricia Piris, Kevin Muller, Guillaume Pinna, Tim W. Failes, Greg M. Arndt, Nathalie Baeza-Kallee, Carole Colin, Olivier Chinot, Diane Braguer, Xavier Morelli, Nicolas André, Manon Carré, Emeline Tabouret, Dominique Figarella-Branger, Marion Le Grand, Eddy Pasquier

## Abstract

Synergistic drug combinations are an attractive anticancer strategy but prove challenging to identify. Here we present a stepwise approach consisting in revealing core cancer vulnerabilities and exploiting them through drug combination screen to uncover synergistic treatments for glioblastoma patients.

**Methods:** We established an innovative method, based on high-throughput screening, target deconvolution and functional genomics, to reveal core vulnerabilities in glioblastoma. Combination drug screen targeting these vulnerabilities was then designed to unveil synergistic associations. The therapeutic potential of the top drug combination was validated in two different clinically-relevant models: an organotypic *ex vivo* model and a syngeneic orthotopic mouse model of glioblastoma.

**Results:** Large-scale monotherapy drug screening identified 83 potent anti-glioblastoma compounds. Target deconvolution using public chemoinformatic databases uncovered 1,100 targets and interactors of the hit compounds. Screening of a focused siRNA library targeting the top 292 drug interactors revealed 22 targetable vulnerabilities, 9 of which were confirmed as core glioblastoma vulnerabilities by mining the CRISPR screen cohort data from the online Cancer Dependency Map portal. Six selective inhibitors of the core vulnerabilities were then screened in combination with a custom-made library of 88 compounds and synergies amongst the 528 tested pairwise combinations were predicted. The combinations of CHK1 / MEK and AURKA / BET inhibitors were highlighted and validated in 3D tumor spheroids. Using an organotypic *ex vivo* model and a syngeneic orthotopic mouse model, we definitively ascertained the efficacy of dual AURKA / BET inhibition in glioblastoma.

**Conclusions:** Collectively, we uncovered that dual inhibition of BET proteins and aurora kinase A is highly synergistic against GBM. Moreover, our study indicates that our approach to exploit drug poly-pharmacology for the rational design of drug combination screens represent a valuable strategy to discover synergistic treatments against refractory cancers.

## INTRODUCTION

Next generation sequencing has revolutionized our understanding of the molecular pathways that drive the development and progression of human cancers [1]. The establishment and expansion of large-scale cancer databases have allowed scientists to mine and analyze an unprecedented number of patient-derived datasets that could lead to the identification of novel targetable vulnerabilities [2]. Nonetheless, translation of these major findings into clinical practice has been slower than expected. This could be partly explained by the current dominant paradigm in drug discovery based on the identification of one specific disease-related target leading to the design of selective inhibitors/activators of this target. However, it is now well-established that most effective drugs act on multiple rather than single targets and combination therapy is the most effective strategy to treat complex diseases, such as cancer [3,4]. In this context, drug poly-pharmacology could open major therapeutic avenues [5-7]. The number of possible combinations being almost infinite, data-driven approaches to find optimal treatments are needed. Here, we hypothesized that screening existing drugs (either already-approved or in clinical development) associated with target deconvolution represent a unique opportunity to exploit drug poly-pharmacology for the identification of core vulnerabilities in cancer. Those targets can then be used to rationally design biology-driven drug combination screens to find synergistic treatments.

Gliomas are the most common primary brain tumors in neuro-oncology practice of adults. About half of all newly diagnosed gliomas are classified as glioblastoma (GBM), which is the most malignant type of brain cancer defined as grade 4 in the 2021 WHO Classification [8]. More than 90% of all cases are classified as primary tumors that arise *de novo*. GBM were previously classified into four subtypes based on transcriptional features: classical, proneural, mesenchymal and neural. However, integrating the malignant cell programs, their plasticity and their modulation by genetic drivers as well as certain molecular markers, including TERT promoter mutation, IDH1/2 mutation, *MGMT* promoter methylation or the combination of whole chromosome 7 gain / 10 loss, demonstrate the high intratumor heterogeneity and can now provide powerful prognostic information [9,10]. Median patient survival is approximately 15 months and less than 7% of patients survive 5 years after diagnosis, contributing to 3–4% of all cancer-related deaths worldwide [11]. Despite recent advances in our understanding of glioma biology, the current standard-of-care for GBM is still based on tumor resection with concurrent radiotherapy and chemotherapy with temozolomide [11]. Nevertheless, in the majority of cases, acquired resistance to both radiation and temozolomide occurs. Thus, there is an urgent need to establish innovative strategies and discover new treatments to improve the outcome of patients with GBM.

In this study, we applied a chemogenomic screen targeting core vulnerabilities in GBM in order to quickly identify synergistic combinations. We uncovered that dual inhibition of BET proteins and aurora kinase A is highly synergistic against GBM using 3D spheroids, organotypic *ex vivo* and syngeneic orthotopic *in vivo* mouse models. Altogether, our study indicates that exploiting core vulnerabilities to design a biology-driven drug screening represent a valuable strategy to discover highly synergistic drug combinations against refractory cancers.

## METHODS

### Cell culture

The human U87 (RRID: CVCL_0022), U87vIII and T98G (RRID: CVCL_0556) GBM cell lines were kindly provided by the Children’s Cancer Institute (Sydney, Australia) while the murine GL261 (RRID: CVCL_Y003) GBM cell line was obtained from ATCC. They were grown in Dulbecco’s Modified Eagle Medium (Gibco, #11960044) containing 10% Fetal Calf Serum, 1% sodium pyruvate (Gibco, #11360070) and 1% penicillin-streptomycin (Gibco, #15140122). Stable U87 and GL261 cell lines were established by transfection of an mtDsRed plasmid with Lipofectamine 2000 (Invitrogen, #11668019) according to the manufacturer’s protocol, followed by geneticin selection (0.8mg/mL, Gibco, #10131035) and two cycles of fluorescence-activated cell sorting. All cell lines were regularly screened and are free from mycoplasma contamination. Cells were incubated in a humid atmosphere at 37°C with 5% CO2.

### Drugs and reagents

Alisertib (10mM, Selleckchem. #s1133), Birabresib (10mM, Selleckchem. #s7360), Prexasertib (10mM Selleckchem. #s6385), Mirdametinib (10mM Selleckchem. #s1036), and Panobinostat (10mM Selleckchem. #s1030) were resuspended in dimethyl sulfoxide (DMSO, Sigma, #D5879). Stock solutions were stored at -20°C. The solutions were diluted in culture medium extemporaneously for the experiments.

### High-throughput drug screening

Three libraries (Prestwick chemical, Lopac and Tocris) containing ∼2,800 unique FDA-approved drugs and pharmacologically-active molecules were screened in a primary screen on U87 cells at a single dose of 5μM. Cell viability was assessed after 72h incubation using a home-made Alamar Blue solution (75mg resazurin, 12.5mg methylene blue, 164.5mg potassium hexacyanoferrate III and 211mg potassium hexacyanoferrate II trihydrate, 500ml water; Sigma) as previously described [12]. Assay performance was evaluated by calculating the Z-factor for each assay plate, according to the method described by Zhang *et al* [13]. Compounds were considered candidates when they significantly inhibited cell viability by at least 50%. The top 280 candidates were then screened in U87, U87vIII and T98G cell lines, using the same methodology. Compounds were defined as confirmed hits when they significantly inhibited cell viability by at least 50% in 2 out of the 3 tested GBM cell lines.

### Focused siRNA library generation

All the known targets and interactors of the pharmacological hits were listed by interrogating three complementary databases: DrugBank (https://go.drugbank.com/), Gostar (https://www.gostardb.com/gostar/) and DRUGSURV (http://www.bioprofiling.de/GEO/DRUGSURV/) [14]. This yielded a total of 1,100 known targets and interactors, of which 292 were associated with at least 3 pharmacological hits. A focused siRNA library (Silencer™ Select pre-designed siRNA; ThermoFisher Scientific) was designed to individually silence the genes encoding the 292 targets / interactors, with 3 different and specific siRNA sequences per gene.

### High-throughput siRNA screening

For screening purpose, an automated reverse transfection protocol was developed on a robotic workstation equipped with a 96-well head probe (Nimbus, Hamilton). Each siRNA sequence from the library was transfected as a separate triplicate in different well positions of 3 independent culture plates to minimize positional errors. Each culture plate also received different positive and negative controls. Eight wells received the transfection reagent alone (“MOCK” well, negative controls), 16 were transfected with a scrambled siRNA (“NEG” wells, negative control, Ambion), and 8 were transfected with a pool of cytotoxic siRNAs (“AllStars” wells, positive control, Allstars maximal death control, Qiagen). Briefly, siRNA sequences were lipoplexed with Lipofectamine RNAiMAX (Life Technologies, #13778150) in clear bottom, black-walled 384-well culture plates (Greiner μClear plates, #781091). After 15min of complexation, GBM cell lines (U87, U87vIII and T98G) were seeded on top of the lipoplexes (1,000 cells/well; final [siRNA] = 5nM) and incubated for 3 days at 37°C and 5% CO2 in a humidified incubator. Cells were then fixed with Paraformaldehyde 4% (Sigma, #1004969011), stained with Hoechst 33342 and plates were imaged on a High-Content Imaging device (Operetta HCS epifluorescence microscope, Perkin Elmer). Three fields per well were acquired at 10x magnification in blue channel (excitation: 380 ± 20nm; emission: 445 ± 35nm). Hoechst-stained nuclear regions of interest were segmented using Harmony software (Perkin Elmer) and cell viability was expressed in each plate as the relative number of cellular events in sample wells relative to the average of NEG wells. Genes were considered as candidates when at least 2 out of 3 siRNA sequences gave statistically significant results in at least 2 GBM cell lines (Z-score scoring). Candidate genes were then retested in an identical setup, and genes were accepted as hits when they passed the selection criteria (2/3 active siRNA sequences) a second time and leading to at least 20% of decrease in cell viability in all tested cell lines.

### DepMap database analysis

Using the Dependency Map portal (https://depmap.org/portal/), we extracted from the CRISPR DepMap 22Q2 Public+Score, Chronos dataset, the gene effect scores derived from CRISPR knockout screens published by Broad’s Achilles and Sanger’s SCORE projects. Gene effect scores were inferenced by Chronos (https://genomebiology.biomedcentral.com) and integration of the Broad and Sanger datasets was performed as described in Pacini *et al* [15], except that quantile normalization was not performed. Negative scores imply cell growth inhibition and/or death following gene knockout. Scores are normalized such that non-essential genes have a median score of 0. As a criterion, genes were considered hits when their gene effect scores are < -0.5 in all 3 selected GBM cell lines (U87MG, U251 and T98G – DepMap id: ACH-000075, ACH-000232 and ACH-000571 respectively).

### Functional validation assay

U87vIII and mtDsRed-expressing U87 cells were seeded in T25 cell culture flasks and transfected with 1ml of Opti-MEM medium (Gibo, #11058021) containing 1% of lipofectamine RNAiMax (Life Technologies, #13778150) and 5nM of siRNA. Three different siRNA sequences targeting *RRM1 (*Silencer^®^ Select #s12357, #s12358, #s12359; ThermoFisher Scientific) were used as well as a non-targeting negative control siRNA with no significant sequence homology to mouse, rat or human gene sequences (Silencer^®^ Select #AM4635). Two days later, cells were seeded in 96-well U bottom and low-binding plates in DMEM medium containing methylcellulose at 0.6g/L. Spheroid growth was then evaluated either daily for one week by measuring the fluorescence ratio (575nm: excitation wavelength / 620nm: emission wavelength) in the case of mtDsRed expressing U87 cells or after 8 days by Alamar Blue in the case of U87vIII cells. Both measurements were performed with a PHERAstar^®^ plate reader (BMG). To evaluate the level of gene knock-down, cells were harvested 48h after transfection and total RNA was extracted using RNeasy Plus mini kit (Qiagen, #74136), following the manufacturer’s instructions. Reverse transcription was performed with Onescript^®^ cDNA synthesis kit (Abm, #G236) and qRT-PCR was undertaken using SsoAdvanced Universal SYBR^®^ Green Supermix (Bio-Rad) and a CFX96TM Real-Time System device (Bio-Rad). Gene expression levels of *RRM1* were determined using the ΔΔ*C*t method, normalized to the *YWHAZ* or *GAPDH* control genes. The following predesigned KiCqStart SYBR^®^ Green primers (Merck, Fontenay-sous-bois, France) were used:

- *RRM1* (forward: 5’-CATTGGAATTGGGGTACAAG; reverse: 5’-AATTCCTTTGCTAACTGGAG),
- *GAPDH* (forward: 5’-ACAGTTGCCATGTAGACC reverse: 5’-TTTTTGGTTGAGCACAGG) and
- *YWHAZ* (forward:5’-AACTTGACATTGTGGACATC; reverse: 5’AAAACTATTTGTGGGACAGC).

To evaluate the level of protein knockdown, cells were lysed in RIPA buffer (Tris-HCl 50mM pH 8.0, NaCl 250mM, Triton-X100 0.1%) freshly supplemented with a cocktail of proteases and phosphatases inhibitors (Sigma-Aldrich, #PPC2020), 72h after siRNA transfection. Protein concentrations were determined using the Bio-Rad Protein Assay (Bio-Rad, #5000001). Proteins were separated by SDS-PAGE and electro-transferred onto a nitrocellulose membrane. Primary antibodies were directed against RRM1 (clone EPR8483; Abcam) and α-tubulin (clone DM1A; Sigma Aldrich). Peroxydase-conjugated secondary antibodies (Cell Signaling, #7074 and #7076) and chemiluminescence detection kit (Millipore) were used for visualization with ChemiDoc Touch Imaging System (BioRad). Membranes were probed again with α-tubulin antibody following a step of stripping using a solution of 5% trichloroacetic acid.

### Gene expression analysis on patient samples

Gene expression analysis was conducted using the R2 microarray analysis and visualization platform (http://r2.amc.nl). RNA-seq data were extracted from two independent cohorts providing open access to data acquired from various forms of cancer: the Cancer Genome Atlas (TCGA) database and from normal tissues: the Genotype-Tissue Expression (GTeX) database. GBM TCGA dataset was used and partitioned in five subtypes according to the data available: classical (n=17), mesenchymal (n=27), neural (n=17), proneural (n=24) and not determined (n=455). We used GTeX normal brain tissue data from the following subgroups: caudate (n=246), cortex (n=255), frontal cortex (n=209), nucleus accumbens (n=246), putamen (n=205), spinal cord (n=159) and thalamus (n=202). Median values were recorded using log2 transformation gene expression. Statistical analyses using ANOVA were performed to compare GBM subtypes gene expression to normal brain tissue gene expression. Boxplots were generated using GraphPad Prism 9.4.1.

### Protein expression analysis on patient samples

Ninety-seven patients were included in this study: they had histologically confirmed GBM, *IDH*-wt [16], were between the ages of 18-70, were not included in experimental therapeutic protocols and were homogeneously treated by the Stupp protocol (Temozolomide + radiotherapy). Tissue microarrays (TMA) were constructed from routinely processed formalin-fixed paraffin-embedded tumor material. Areas of viable and representative tumor following review of all blocks were marked by a pathologist (Pr Figarella-Branger) prior to inclusion into the TMA (3 × 0.6mm cores for each tumor). Immunohistochemistry study was performed on TMA. After steam-heat induced antigen retrieval, 5μm sections of formalin-fixed paraffin-embedded samples were tested for the presence of RRM1 (Rabbit monoclonal, clone EPR8483; Abcam). A Benchmark Ventana autostainer (Ventana Medical Systems SA, Illkirch, France) was used for detection and TMA slides were simultaneously immunostained in order to avoid inter-manipulation variability. Immunostaining was scored by a pathologist (Pr Figarella-Branger), analysing for each triplicate the core demonstrating the strongest immunoreaction, and using the immunoreactive score (IRS) previously described by Casar-Borota *et al* [17]; the IRS (0–12) being the product of the proportion of immunoreactive cells (0, 0%; 1, < 10%; 2, 10-50%; 3, 51-80% or 4, > 80%) and the staining intensity (0, no staining; 1, weak; 2, moderate and 3, strong). The Kaplan–Meier method was used to estimate survival distributions. Log-rank tests were used for univariate comparisons. Cox proportional hazards models were used for multivariate analyses and for estimating hazard ratios in survival regression models. Multivariate analysis included extent of surgical resection. All the tests were two-sided and *p* < 0.05 was considered significant for each statistical analysis. Statistical analyses were conducted using the PASW Statistics version 17 (IBM SPSS Inc., Chicago, IL, USA).

### Drug combination screening

Briefly, 4,000 living cells from the GL261 murine GBM cell line were seeded per well with the Certus Flex^®^ (GyGer) in 384-well plates (Corning, #3830). Cells were incubated in the presence of a custom-made drug library containing 88 drugs alone (see supplementary Table S7 for details) or in association with Alisertib 1μM, Prexasertib 10nM, Vistusertib 250nM, Dinaciclib 5nM, Rigosertib 250nM or Panobinostat 10nM. Drugs were distributed with the Echo 550 liquid dispenser^®^ (Labcyte) at 6 different concentrations covering 3 logs (*i*.*e*., 1nM to 1μM, 10nM to 10μM or 100nM to 100μM) in constant DMSO. Cell viability was measured using CellTiter-Glo^®^ 2.0 Cell Viability Assay (Promega, #G9243) after 72h of drug incubation in a humidified environment at 37°C and 5% CO2. Luminescence was measured using a PHERAstar^®^ plate reader (BMG). Data were normalized to negative control wells (DMSO only). IC50, defined as half maximal inhibitory concentration values, and AUC (Area Under the Curve – %.mol.L^-1^) were obtained using library(ic50), library(drc), library(ggplot2) and library(PharmacoGx) packages from R studio.

### 3D spheroid cell viability assay

The mtDsRed-expressing U87 and GL261 GBM cells were seeded at 2,000 and 4,000 cells/well, respectively in 96-well U bottom and low-binding plates in DMEM medium containing methylcellulose (0.3g/L). After 24h and 6 days, cells were treated with a 6×5 combination matrix containing two compounds at different concentration ratios. After 15 days of drug treatment, metabolic activity was detected by addition of Alamar blue and spectrophotometric analysis using a PHERAstar^®^ plate reader (BMG). Cell viability was determined and expressed as a percentage of untreated control cells. To assess the synergy score of the tested combinatorial treatments, Bliss heatmaps were generated using the SynergyFinder web application (https://synergyfinder.fimm.fi) [18].

### Microscope image acquisition

Images of U87vIII and mtDsRed-expressing U87 spheroids were acquired by phase contrast and RFP fluorescence channel using a 4x/0.13 objective with an EVOS FL^®^ Cell Imaging System (Invitrogen) and images of mtDsRed-expressing U87 and GL261 spheroids were acquired by RFP fluorescence channel using a 4x/0.1 objective with a Leica DM IL LED^®^ (Leica) at the end point of experiments.

### Brain organotypic model development and analysis

To establish organotypic cultures, the healthy brains of Swiss immunocompetent mice over 12 weeks old were surgically harvested and sectioned into 250μm thick slices using a vibrating blade microtome (Vibratome 2000s, Leica Biosystems). A 4-days spheroid formed from 4,000 mtDsRed-expressing GL261 GBM cells was then grafted onto each brain slice. These organotypic co-culture models were then placed on inserts with 0.4μm pore size membranes (Falcon®, #353090) and maintained in medium containing 50% MEMa, 25% horse serum (Gibco, #16050122), 25% Hanks’ Balanced Salt Solution (HBSS; Gibco, #14065056), 10mM HEPES buffer (Gibco, #15630106), 28mM Glucose (Gibco, #15023021), 1% L-Glutamine (Gibco, #25030081) and 1% penicillin-streptomycin (Gibco, #15140122). After daily exposure to 2.5μM of Alisertib, 1μM of Birabresib or their combination for 5 consecutive days, tumor growth and invasion within the brain slices were analysed over time, using the JuLI™ Stage imaging system and the PHERAstar^®^ plate reader (BMG) (λex 540nm/λem 580nm – fluorescence signal acquisition with a 30×30 matrix scanning mode).

### Animal study

Six-week-old C57BL/6 mice were anesthetized and 50,000 murine GL261 GBM cells were injected in the corpus callosum (1mm anterior to bregma, – 1mm lateral and – 2mm of the cortex surface) using a stereotaxic frame (Kopf). Animals were observed until they fully recovered. A total of 40 mice were included in the study and randomized into 4 groups (n > 9) one week after tumor cell injection. Mice thus received oral administrations of Alisertib (25mg/kg in 10ml/kg application volume), Birabresib (75mg/kg in 10ml/kg application volume), Alisertib + Birabresib (25mg/kg + 75mg/kg in 10ml/kg application volume), or vehicle only (10ml/kg, vehicle groups) from day 7 to 28 after GBM implantation (5 administrations / week). Alisertib and Birabresib were resuspended in a mixture of DMSO (10%) and corn oil (90% – Sigma) to ensure full dissolution and achieve better gastrointestinal absorption of the drugs. The body weight and general status of mice were recorded every day. Mice were euthanized when they lost more than 20% of their initial body weight or displayed signs of distress or neurological deficits. All animal experiments were performed by adequately trained research personnel and were authorized by the local animal ethics committee (authorization number #02201902).

### Histology and immunohistochemical analysis

Serial 4-μm paraffin sections of brains were stained with hematoxylin and eosin and examined under the microscope for the presence of tumors. Immunohistochemistry was performed on adjacent paraffin sections with monoclonal anti-Ki67 antibody (clone MIB-1; Dako) and avidin–biotin–peroxidase method (*Vectastain* Elite *ABC* kit, *Vector* Laboratories).

### Statistical analyses

Each experiment was performed at least in triplicate. Data are presented as mean ± SD or S.E.M as indicated in the figure legends. Statistical significance was tested using unpaired Student’s t test. For experiments using multiple variables, statistical significance was assessed *via* two-way ANOVA. Log-rank (Mantel-Cox) and Gehan-Breslow-Wilcoxon tests were used to compare animal survival by Kaplan-Meier analysis. A significant difference between two conditions was recorded for \**p* < 0.05; \*\**p* < 0.01; \*\*\**p* < 0.001.

## RESULTS

### Stepwise chemogenomic screen identifies nine core vulnerabilities in GBM

To first unveil new disease-relevant targets through the exploitation of drug poly-pharmacology in GBM, we performed a high-throughput drug screen with three commercially available libraries containing small molecules that have well-annotated pharmacology and are suitable for phenotypic screens. More than 2,800 unique already-approved drugs and pharmacologically active molecules (Table S1) were screened in the U87 GBM cell line and the top 280 primary candidates leading to a decrease of more than 50% of cell survival were selected and used in a secondary screen with 3 different GBM cell lines (Table S2). Amongst the primary hits, 83 reduced the cell viability of at least 2 out of the 3 tested cell lines by more than 50% and were further confirmed as “GBM killers”. Besides experimental drugs, which account for ∼41% of the pharmacological hits (34/83), the high-throughput drug screen also identified 19 chemotherapy agents (∼23%), 3 targeted therapies and 27 non-oncology drugs, out of which 15 are FDA-approved drugs currently used for human diseases and that could potentially be repurposed for the treatment of GBM (Figures 1A-B and Table S3 for details). As the second step of our methodology, we performed target deconvolution using three chemoinformatic databases (DrugBank, Drugsurv and Gostar) to exploit drug poly-pharmacology. All the known molecular targets and interactors of the 83 pharmacological hits were listed (Table S4). Amongst the 1,100 identified interactors, 292 were associated with at least three pharmacological hits and were used to design a focused siRNA library containing three individual sequences for each of the 292 top targets. As a third step, siRNA screening was performed in three GBM cell lines, resulting in the identification of 22 pharmacologically targetable hits that decreased by at least 20% the viability of all tested GBM cell lines (Figure 1C and Table S5 for details). Finally, to prioritize targets, we cross-evaluated our siRNA screening results with the CRISPR loss-of-function screen data cohort from the Cancer Dependency Map Project (https://deepmap.org). Nine of the 22 genes identified by our focused siRNA screen were defined as essential genes (gene effect score < -0.5) in all 3 tested GBM cell lines: *RRM1, PLK1, CHEK1, AURKB, CDK1, AURKA, FRAP1, HDAC3* and *ATP1A1* (Figure 1D and Table S6).

**Figure 1.**
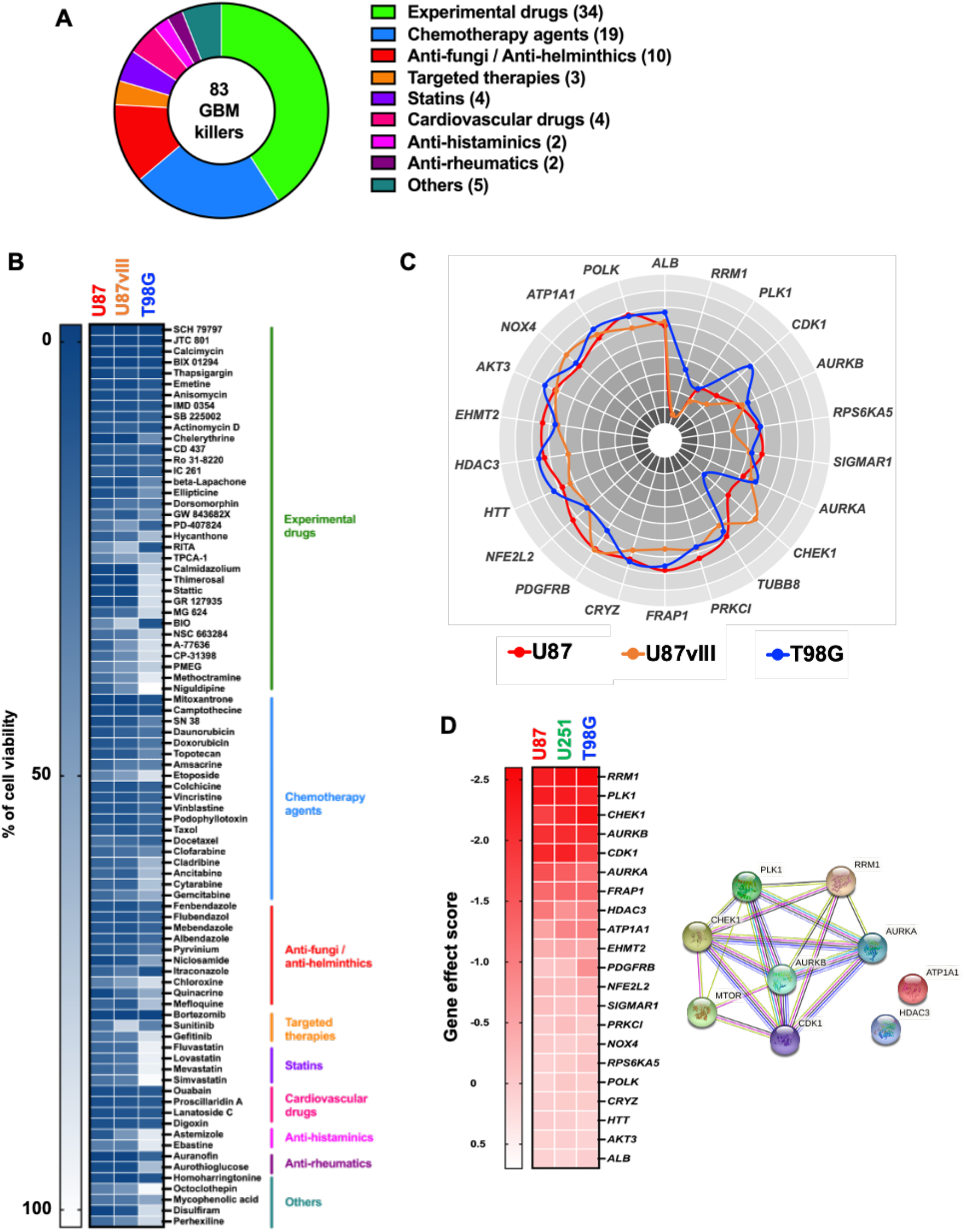
Stepwise chemogenomic screen identifies nine actionable gene vulnerabilities in GBM. (**A**) Primary screen was performed in U87 GBM cell line with more than 2,800 unique compounds tested alone (5μM). After 72h incubation, cell viability was assessed using Alamar Blue. Top 280 primary candidates were then re-tested in a secondary validation screen in U87, U87vIII and T98G GBM cell lines using the same protocol. Eighty-three compounds were defined as GBM killers in at least 2 GBM cell lines and are represented in donut diagram and classified by pharmacological classes. (**B**) Heat map classification representing the cell viability in all tested GBM cells of the 83 hit compounds. (**C**) By target deconvolution using pharmacological online databases, 1,100 known targets and interactors of the 83 hit compounds were revealed. Amongst them, 292 were targeted by at least 3 hit compounds and selected to build a focused siRNA library. Three GBM cell lines (U87, U87vIII and T98G) were individually transfected with 3 siRNA sequences (5nM) for each of the 292 targets / interactors. Cell viability was assessed by high-content imaging following Hoechst 33342 staining. Polar plot of 22 gene hits, which decreased the cell viability in each tested cell lines by at least 20%. Polar plot was made up of 10 data rings, each radial point representing a ten percent increment of the cell viability on a scale from 0 (inner radial point) to 100 (outer radial point). (**D**) Heat map representing in 3 GBM cell lines the gene effect score of the 22 gene hits extracted from the CRISPR screen cohort data from the online Dependency Map portal. Right panel shows the protein-protein interaction network of the 9 hits identified as core vulnerabilities in all tested GBM cell lines (Gene effect score < -0.5; https://string-db.org/).

To ascertain the potential of our methodology to reveal core vulnerabilities in GBM, we focused our functional study on the top gene hit *RRM1*, which encodes for the catalytic subunit of an enzyme playing a key role in the production of deoxyribonucleotides (dNTPs) for DNA replication [19]. A transient knockdown using three different siRNA sequences leading to a significant inhibition of RRM1 gene and protein expression (Figures 2A-B) resulted in a decrease in tumor spheroid growth, observed at day 5 and maintained until day 8. This was reflected by a significant drop in the U87 tumor spheroid growth of 70 +/-10%, 56 +/-16% and 72 +/-6% for the siRNA sequences #1, #2 and #3 in comparison to control, respectively (*p* < 0.001, Figure 2C). Similar results were obtained with the U87vIII GBM cell line, in which *RRM1* knockdown resulted in a 65-72% decrease in tumor spheroid viability after 8 days in comparison to control (*p* < 0.001, Figures S1A-C). Our transcriptomic analysis using freely available datasets in the R2 platform showed that *RRM1* expression is significantly up-regulated in GBM subtypes as compared to normal brain tissue (8.2 +/-0.2 and 3.3 +/-0.3, respectively; p < 0.0001; Figures S1D). Furthermore, we evaluated RRM1 protein expression in GBM patient samples using immunohistochemistry staining of tissue microarray (Figure 2D). On the 97 analysed GBM samples, 69 cases (71%) were positive for RRM1 expression (24 cases were negative and 3 cases were not informative). We found that high expression of RRM1 (IRS from 4 to 12) was significantly associated with a poor outcome in univariate analysis (p < 0.01) and it remained prognostic in multivariate analysis (adjusted by type of surgery) (p=0.003; HR=1.991, 95% CI [1.259 – 3.147]). Altogether, these results demonstrate that our stepwise chemogenomic screening approach is an efficient strategy to exploit drug poly-pharmacology for the identification of core vulnerabilities, that could be further therapeutically exploited.

**Figure 2.**
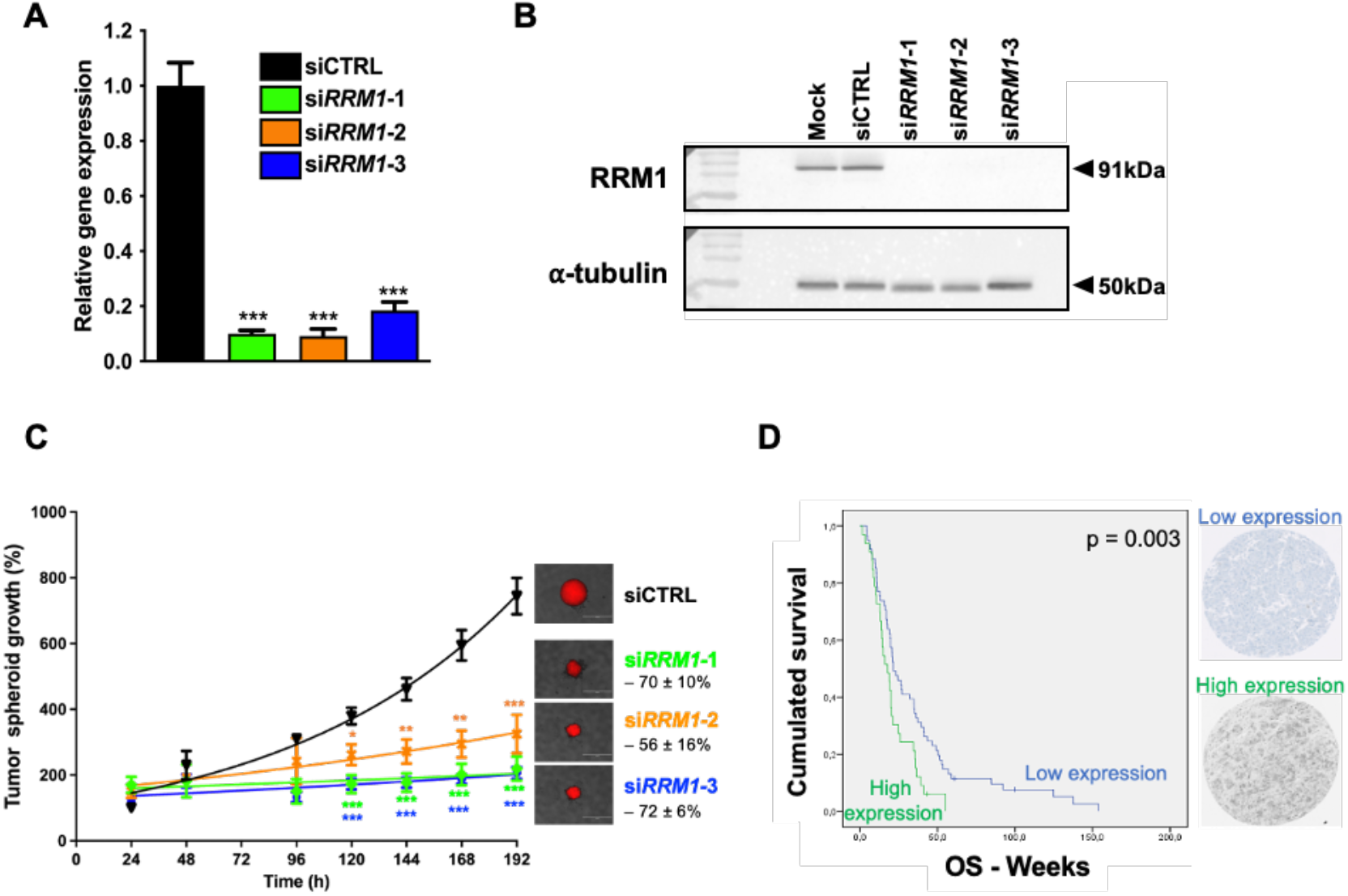
Functional validation of top gene hit *RRM1* in glioblastoma cells and patient samples. (**A**) *RRM1* relative gene expression following 48h transfection of U87 cells with negative control siRNA and 3 different siRNA sequences targeting *RRM1*, as evaluated by qRT-PCR using *YWHAZ* as housekeeping gene. *Bars*, mean of at least 4 independent experiments; *Error bars*, S.E.M; ***, p<0.001. (**B**) Representative western blot showing RRM1 protein expression following 72h siRNA transfection, using α-tubulin as loading control. (**C**) The mtDsRed-expressing U87 tumor spheroid growth following siRNA transfection assessed by daily fluorescence measurements. *Points*, mean of at least 4 independent experiments; *Error bars*, S.E.M; *, p<0.05; **, p<0.01; ***, p<0.001. Representative photographs of tumor spheroids at day 8 and mean decrease in spheroid growth +/-S.D are included in insert (*right*). *Scale bar*, 1mm. (**D**) Kaplan-Meier survival estimate of GBM patients according to RRM1 protein expression assessed by immunohistochemistry. Cox proportional hazards regression *p* value is shown and representative pictures of tumor samples displaying low (*top*) and high (*bottom*) RRM1 expression are included (*right*).

### Targeting core vulnerabilities uncovers synergistic drug combinations in GBM

In order to therapeutically exploit the core vulnerabilities identified by our chemogenomic screens, we established a biology-guided drug combination screen using available inhibitors of 6 core vulnerabilities: AURKA inhibitor Alisertib, CDK1 inhibitor Dinaciclib, HDAC3 inhibitor Panobinostat, CHEK1 inhibitor Prexasertib, PLK1 inhibitor Rigosertib and mTOR inhibitor Vistusertib (Figure S2A). These inhibitors were screened in the murine GL261 GBM cell line in combination with a custom-made library of 88 FDA-approved compounds, of which 15 repurposed drugs identified as pharmacological hits in our first drug screening (Figures 1A&B) together with 15 epidrugs, 49 targeted therapies and 9 additional repurposed drugs. These agents were chosen because they were either approved by the US Federal Drug Administration (FDA) or in advanced clinical development and were considered potentially useful in treating brain cancer patients based on prior clinical and preclinical evidence (Table S7). We used the difference of area under the dose-response curve (AUC) between the combinatorial treatment and the monotherapy condition. Differences in AUC of more than 5%.mol.L^-1^ were considered as potentially synergistic and below -5%.mol.L^-1^ as potentially antagonistic combinations. A full report of this combination screen can be found in Table S7. Amongst the 528 tested pair-wise combinations, 4.9% appeared to have a potential synergistic effect (Figures S2B-G). Since combinations with Alisertib, Prexasertib, Panobinostat and Vistusertib were the most favorable (4.5, 5.7, 5.7 and 10.2% of potential synergisms, respectively), we focused on these 4 combinatorial treatments (Figures S2D-G). Amongst the 88 tested drugs, 15 were epidrugs including 7 BET inhibitors, 4 HDAC inhibitors and 4 other epidrugs (Figure 3A). Our results indicated that the combination of Alisertib or Vistusertib with most of the BET inhibitors was potentially synergistic, while it was mostly antagonistic when combined with Panobinostat, and to a lesser extent with Prexasertib (Figure 3A). These results were reflected by a decrease in the IC50 of BET inhibitors when co-administered with Alisertib and Vistusertib and, in contrast, an increase in the IC50 of these drugs when co-administered with Prexasertib and Panobinostat (Figure S3A). Regarding the 24 repurposed drug tested in our library, 2 potentially synergistic combinations were observed between Vistusertib / Aripiprazole and Panobinostat / Quinacrine, which was illustrated by a decrease in the IC50 of both repurposed drugs when combined with an mTOR or HDAC3 inhibitor (Figures 3B and Figure S3B). Finally, 49 targeted therapies were tested covering major signaling pathways involved in cancer (Figure 3C). Our combination drug screen highlighted a potential synergism between MEK inhibitors and Alisertib or Prexasertib, while a potential antagonism was observed with Panobinostat and to a less extend with Vistusertib (Figure 3C). This was also demonstrated by the decrease in IC50 values of both MEK inhibitors, Mirdametinib and Selumetinib, when combined with Alisertib or Prexasertib and their increase when associated with Panobinostat or Vistusertib (Figure S3C).

**Figure 3.**
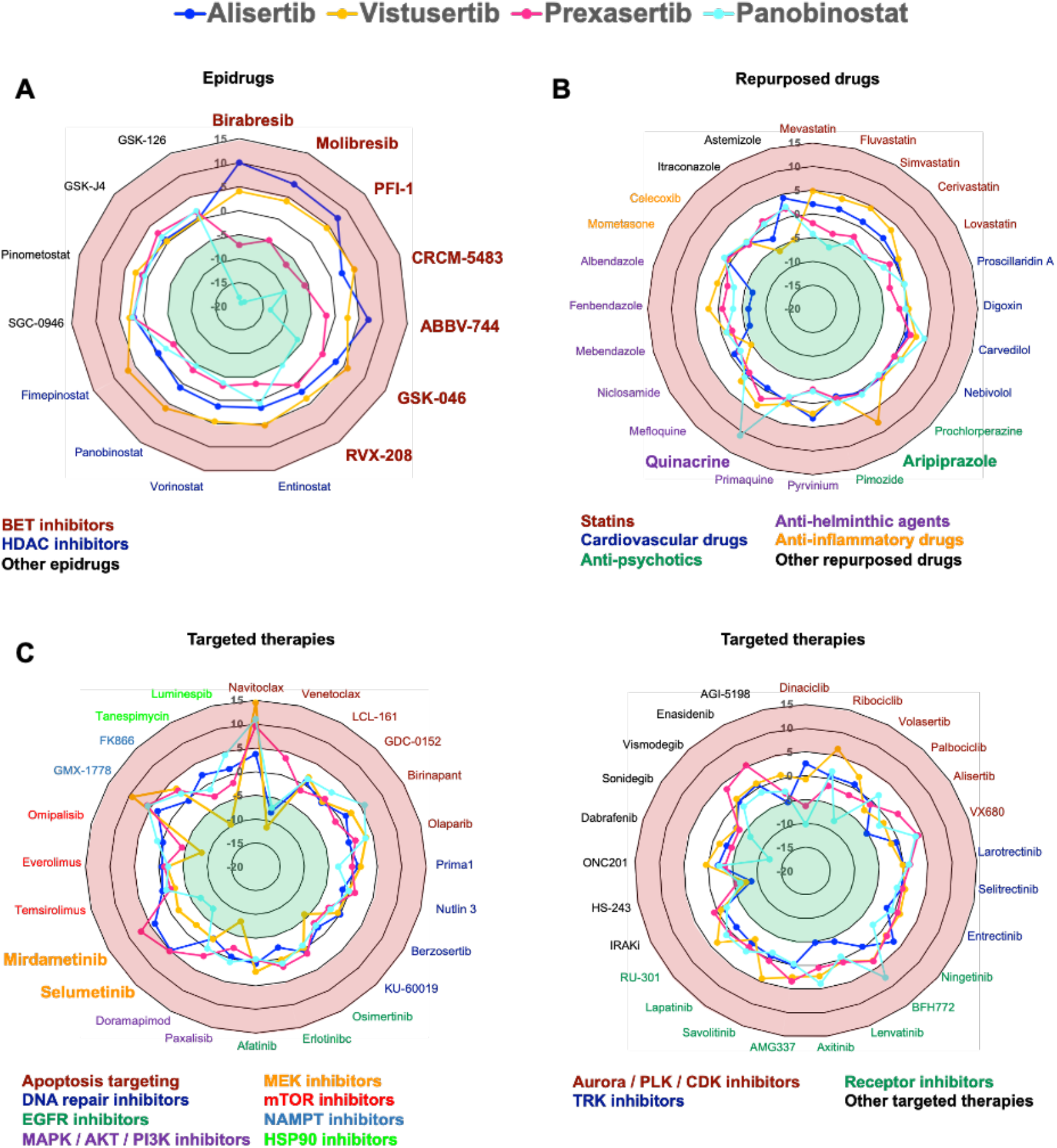
A biology-guided drug combination screen reveals potential synergistic treatments in GBM. A custom-made library containing 88 drugs was screened on the murine GL261 GBM cell line in a dose-effect manner alone or in association with Alisertib 1μM, Vistusertib 250nM, Prexasertib 10nM or Panobinostat 10nM. After 72h of drug incubation, cell viability was assessed using CellTiter Glo^®^. Radar plots show the difference of AUC between the combinatorial treatment and the monotherapy condition (combination with Alisertib: dark blue line, Vistusertib: yellow line, Prexasertib: pink line and Panobinostat: light blue line). Radar plots were made up of 7 data rings on a scale from -20%.mol.L^-1^ (inner ring) to 15%.mol.L^-1^ (outer ring) with an increment of 5%.mol.L^-1^. Green rings represent antagonist combinations (AUC differences < -5%.mol.L^-1^) and the red ones indicate synergistic treatments (AUC differences > 5%.mol.L^-1^). Radar plots representation of data obtained with (**A**) 15 epidrugs, (**B**) 24 repurposed drugs, and (**C**) 49 targeted therapies.

To confirm the interactions underlined by our biology-driven drug combination screen, we assessed them using the human U87 and murine GL261 GBM cell lines in 3D spheroid culture conditions treated during 15 days. We focused our validation experiments on two potentially synergistic drug combinations: Alisertib / Birabresib (Figures 3A and Figure S3A), Prexasertib / Mirdametinib (Figures 3C and Figure S3C) and one potentially antagonistic combination Panobinostat / Birabresib (Figures 3A and Figure S3A). A 6×5 matrix that contained the two compounds at different concentration ratios was created, allowing to cover a large range of drug combinations. To assess the synergy of the two compound treatments, the Bliss score was calculated using the SynergyFinder web application (https://synergyfinder.fimm.fi) [18]. We found that the combination of AURKA inhibitor Alisertib and BET inhibitor Birabresib produced a highly synergistic effect with a Bliss score of 14.9 +/-1.9 in GL261 and 6.6 +/-0.7 in U87 (Figures 4A&G). Indeed, while a drop in the GL261 spheroid viability was observed with monotherapies of 25nM Alisertib and 100nM Birabresib (16.2 +/-4.1% and 2.7 +/-3.7% respectively, Figures 4D), the synergistic score of their combination was reflected by a decrease in the GL261 spheroid viability of 56.6 +/-3.4% in comparison to untreated cells (Figure 4D). Similar results were obtained with the U87 GBM cell line, in which the association of 25nM Alisertib and 1μM Birabresib resulted in a decrease in tumor spheroid viability of 59.2+/-12.5% after 15 days in comparison to control, while each monotherapy reduced the spheroid viability of 19.8 +/-3.7% and 29.3 +/-3.7% respectively (Figure 4J). Our results also confirmed the synergism between CHK1 inhibitor Prexasertib and MEK inhibitor Mirdametinib, which was also ascertained in the U87 3D spheroid models (Figures 4B, E, H&K). These results were illustrated by a Bliss score of 7.9 +/-3.7 in GL261 and 2.9 +/-2.9 in U87 (Figures 4B&H). Finally, our screen results highlighted potentially antagonistic interactions between BET inhibitors and Panobinostat in GL261 cells (Figures 3A & Supplementary Figure S3A). These observations were confirmed in the GL261 spheroid model when treated with HDAC inhibitor Panobinostat and BET inhibitor Birabresib, as demonstrated by a Bliss score of -2.0 +/-4.0 (Figures 4C&F). In sharp contrast, a synergistic effect was shown in the U87 spheroid model with a Bliss score of 8.5 +/-4.3 (Figures 4I). Indeed, a decrease in the U87 spheroid viability was observed with monotherapies of 25nM Panobinostat and 1μM Birabresib (52.5 +/-9.3% and 22.9 +/-17.7% respectively, Figures 4I&L), that was even more important following combination treatment (87.3 +/-7.1% in comparison to untreated cells; Figure 4L). Collectively, our results underlined that our approach to rationally design drug combination screening represent a valuable strategy to rapidly discover pharmacological synergisms.

**Figure 4.**
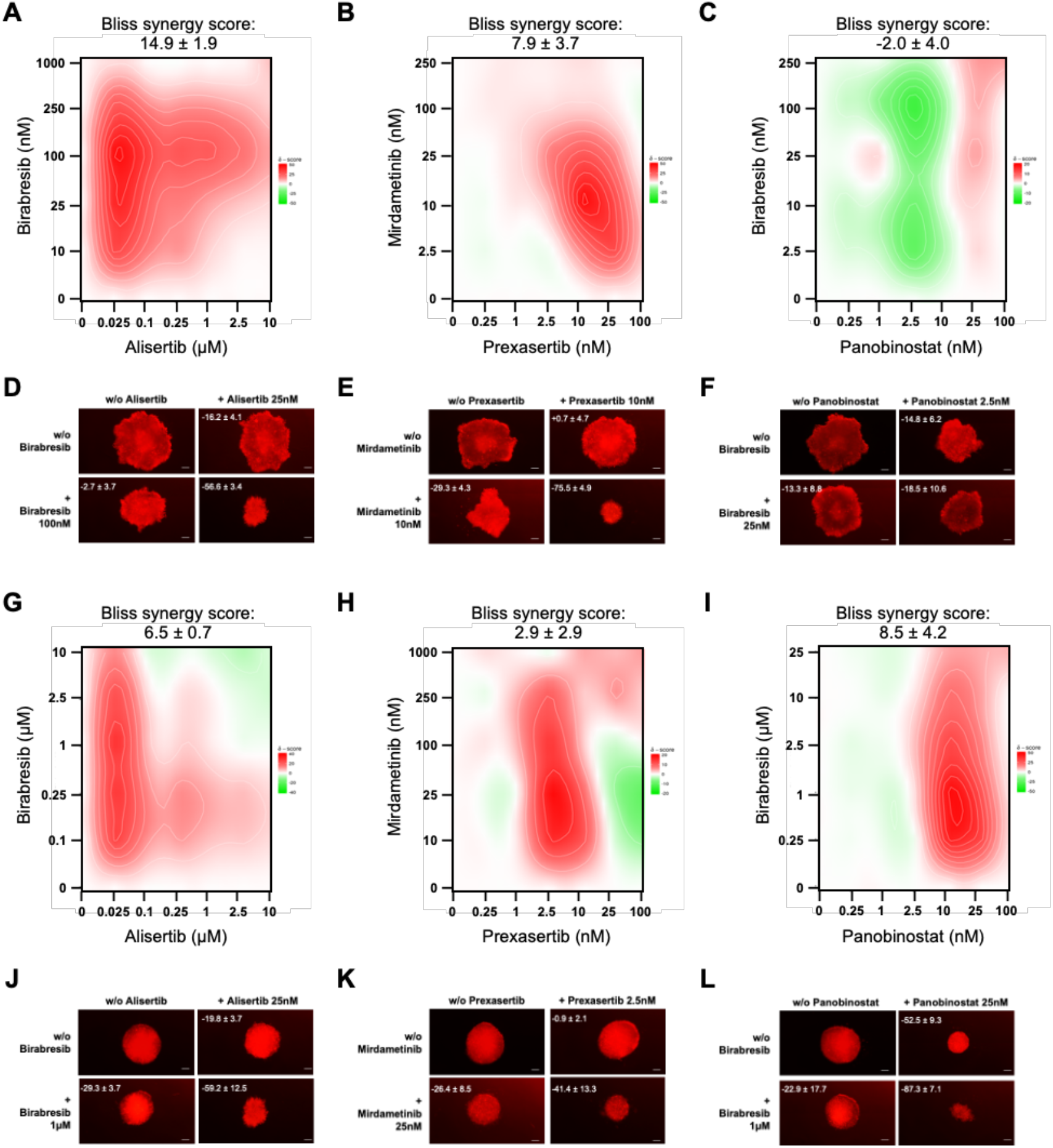
Validation of drug combinations in spheroid GBM models. A 6×5 matrix was used to test drug combinations in GL261 (panels **A** to **F**) and U87 (panels **G** to **L**) spheroid models. Heat maps representing the Bliss score for three tested combinations: Alisertib / Birabresib (panels **A** and **G**), Prexasertib / Mirdametinib (panels **B** and **H**) and Panobinostat / Birabresib (panels **C** and **I**). n= 4. Representative photographs of GL261 (panels **D** to **F**) or U87 (panels **J** to **L**) tumor spheroids at day 14 and mean decrease in spheroid growth +/-S.D are indicated. *Scale bar*, 1mm.

### Dual BET / AURKA inhibition is highly synergistic in GBM models *ex vivo* and *in vivo*

Our data indicated that the Aurora kinase A inhibitor Alisertib was highly synergistic with Birabresib, a pan-BET inhibitor in both tested GBM cell lines (Figures 4A, D, G&J). To further evaluate the potential of dual AURKA / BET inhibition in more clinically-relevant conditions, we developed an organotypic *ex vivo* model in which the murine GL261 GBM spheroids stably expressing mtDsRed were grafted into slices of healthy mouse brain. These innovative cultures were exposed to 2.5μM of Alisertib, 1μM of Birabresib or their combination during 5 consecutive days. After eight days, our data showed that both monotherapies have no significant impact on the growth of GL261 tumor masses (+11 +/-6 and -13 +/-3%, respectively; Figure 5A), while the combinatorial treatment resulted in a significant reduction of 53 +/-2% in comparison to control (p < 0.001; Figure 5A). After 14 days, Alisertib led to a significant decrease in tumor spheroid growth (−36 +/-3.8% in comparison to control; p < 0.05) and so did Birabresib (−58 +/-2.5% in comparison to control; p < 0.01) (Figures 5A-C and S4A). The synergistic effect of Alisertib / Birabresib was even exacerbated over time, with the combination decreasing tumor growth by 81 +/-1.5, 70 +/-2 and 55 +/-1.5% in comparison to control, Alisertib alone and Birabresib alone, respectively (p < 0.0001; Figures 5A-C and S4A). To definitively ascertain the synergism between Alisertib and Birabresib in GBM, we used an orthotopic syngenic mouse model where murine GL261 GBM cells were transplanted by stereotaxy into the corpus callosum of C57BL/6 mice. One week later, mice were randomized and treated *p*.*o*. with either vehicle alone (1/10 DMSO in corn oil), Alisertib alone (25mg/kg), Birabresib alone (75mg/kg) or the combination of both drugs. Monotherapies only marginally extended the median survival from 25.5 days observed for the vehicle control mice to 28 days for Alisertib alone and 30.5 days for Birabresib alone (Figure 5D and Table 1). Our results demonstrated that the combination treatment could significantly increase the median survival of GL261-bearing mice to 48 days (p = 0.0002, 0.0031 and 0.0131 in comparison to vehicle, Alisertib alone and Birabresib alone, respectively; Figure 5D and Table 1). The combination thus increased the therapeutic benefit of Alisertib and Birabresib alone by 9- and 4.5-fold, respectively (Table 1). It is worth to note that two animals treated with the combination were alive and well at study completion (t=120 days). Anatomopathological examination at day 120 revealed a massive tumor mass invading brain parenchyma with high proliferation index in control mice euthanized at day 24 (Figure 5E, right top panel). Conversely, one of the long-term survivors displayed no evidence of disease while the other had a very small tumor with rare proliferating cells, according to Ki-67 immunohistochemistry (Figure 5E, right bottom panel). Moreover, the survival study showed that treatment with Alisertib or Birabresib alone or in combination was well-tolerated and did not result in increased toxicity, as evidenced by a lack of significant change in animal weight during the treatment (Figure S4B). In conclusion, our results indicate that dual inhibition of BET proteins and Aurora kinase A is highly synergistic and could be used as an alternative therapeutic strategy in GBM.

**Table 1:** Median survival of mice harboring GL261 orthotopic xenograft tumors.

| Treatment (mice per cohort) | Median survival (days) | Log-rank test (vs. vehicle) | Log-rank test (vs. Alisertib) | Log-rank test (vs. Birabresib) |
| --- | --- | --- | --- | --- |
| Vehicle (n=10) | 25.5 | N/A | 0.0409 | 0.0045 |
| Alisertib (n=9)<br>25mg/kg per os, 5 days per week | 28 | 0.0409 | N/A | 0.2545 |
| Birabresib (n=10)<br>75mg/kg per os, 5 days per week | 30.5 | 0.0045 | 0.2545 | N/A |
| Alisertib + Birabresib (n=9) | 48 | 0.0002 | 0.0031 | 0.0131 |

**Figure 5.**
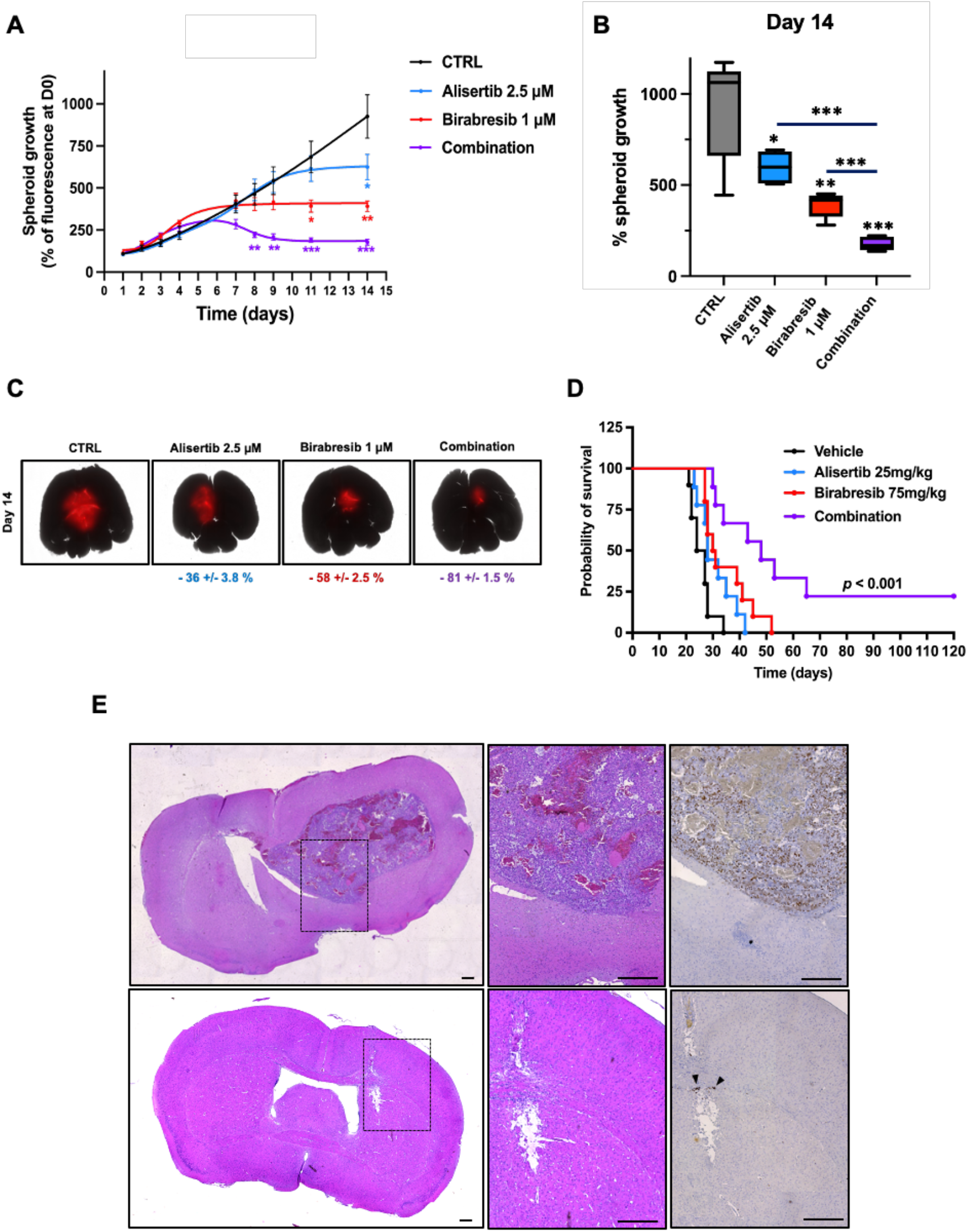
Alisertib / Birabresib combination is highly effective in *ex vivo* GBM organotypic model and in *in vivo* orthotopic mouse model. For five consecutive days, the organotypic brain co-cultures were exposed to daily doses of Alisertib 2.5μM alone (blue), Birabresib 1μM alone (red) or their combination (purple). (**A**) GL261 tumor growth was measured over 14 days by acquisition of the mtDsRed signal with the Pherastar^®^ plate reader (well-scanning mode). Values are the average of n = 5 samples per condition, *Error bars*, S.D; *, p<0.05; **, p<0.01; ***, p<0.001. (**B**) Box plots representing the GL261 tumor growth at the end point. p<0.05; **, p<0.01; ***, p<0.001. (**C**) Representative photographs, acquired with the JuLi™ Stage live imaging system, of mtDsRed-expressing GL261 tumor micro-masses grafted in slices of healthy mouse brain. *Scale bar:* 1mm. Results were expressed as percentage of growth inhibition in treated *vs* control organotypic models (CTRL). S.D. (**D**) Kaplan-Meier survival of murine GBM-bearing mice treated by oral gavage, one week after orthotopic injection of GL261 cells five time a week over three weeks, with vehicle only (1/10 DMSO in corn oil; *black*), Alisertib (75mg/kg; *blue*), Birabresib (25mg/kg; *red*) or the combination of both drugs (*purple*). ***, p<0.001 – log rank test. (**E**) Hematoxylin–eosin and Ki67 staining of GL261 tumors from mice treated with vehicle (a and b) or combination of Alsertib and Birabresib (d and e). Ki67 immunostaining of GL261 tumor treated with vehicle (c) or combination of Alsertib and Birabresib (f). Arrowheads show Ki67 positive cells. Full brain coronal section (a and d) and magnification (b, c, e and f). Scale bar: 500μm.

## DISCUSSION

Despite significant advances in the understanding of GBM biology in recent years, no major breakthrough has been translated into the clinic yet and the standard-of-care has been solely based on radiotherapy with Temozolomide for more than 15 years [8]. In this study, we developed a data-driven approach in order to reveal core vulnerabilities and highlight potential synergistic drug combinations for this unmet medical need.

The predominant paradigm in drug discovery has been the search for maximally selective drugs that act on individual targets. The limitations of this approach is perfectly illustrated by the discovery of EGFR mutation in ∼40% of GBM patients [20], yet EGFR targeted therapies failing in clinical trials due to inefficient drug penetration and distribution in the brain as well as the very high inter- and intra-tumor heterogeneity of GBM [21]. In this context, combination therapy is the most efficient strategy since the association of two drugs can be synergistic and thus allow better anti-tumor response with lower concentration of single agents. Nevertheless, with more than 5,000 approved-drugs or compounds in clinical development, pinpointing synergistic drug combinations represents a major challenge. Herein, we aimed at streamlining the identification of potent combinations by exploiting drug poly-pharmacology using commercially available libraries of compounds with known targets. We hypothesized that the annotated targets of a pharmacological hit may be involved in modulating GBM cell survival, and could thus represent targetable vulnerabilities. We developed a pipeline for the deconvolution of readily druggable targets directly from a high-throughput screening based on drug repurposing principles, followed by a focused siRNA library screen to prioritize targets. As a result, we identified nine core vulnerabilities in GBM: *RRM1, PLK1, CHEK1, AURKB, CDK1, AURKA, FRAP1, HDAC3* and *ATP1A1*. Large pharmacological screens are rarely followed by target validation studies, which prevents the discovery of potential biomarkers. Here to ascertain our method, we focused our functional validation experiments on the top gene hit of our chemogenomic screen approach *RRM1*. Ribonucleotide reductase (RNR) is the key enzyme that catalyzes the production of deoxyribonucleotides (dNTPs) for DNA replication [19]. Several studies have shown that *RRM1* could act as a tumor suppressor gene in lung and pancreas carcinomas [22-24]. These effects have been notably linked to the induction of PTEN expression and an increase in DNA damage repair [25,26]. Moreover, RRM1 is also the predominant cellular determinant of the efficacy of the nucleoside analogue drug gemcitabine. Indeed, studies have reported that high RRM1 expression was associated with a poor prognosis in patients treated with gemcitabine for their non-small cell lung, pancreatic, biliary or bladder cancers [27-30]. Herein, we provide strong evidence supporting an oncogenic role of *RRM1* in GBM. Our results are consistent with recent data demonstrating that a high RRM1 expression leads to GBM growth and poor patient outcome [31.32]. Moreover, RRM1 is highly expressed in brain metastasis and associated with Temozolomide resistance [33,34]. In this study, our results highlight the prognostic role of RRM1 in GBM and advocate for the rapid development of selective RRM1 inhibitors with good brain penetration since current RRM1 inhibitors, including gemcitabine and clofarabine, cannot cross the blood brain barrier. Further studies focusing on associated targets regulating RRM1 oncogenic role in GBM are necessary to fully decipher its functions and explain the observed discrepancy of RRM1 functions between different cancer types.

Multiple high-throughput screening campaigns have been performed to discover new therapeutic options in cancer. The emergence of the CRISPR-Cas9 technology has enabled genome-wide screens, which allow the identification of core vulnerabilities governing cell survival across all cell types, including in GBM [35-37]. However, these functional studies have rarely been combined with pharmacological screens to therapeutically exploit the identified vulnerabilities. Moreover, to reach effective and sustained clinical responses, drug combinations that simultaneously inhibit multiple pathways in cancer cells are needed. Most studies in the field usually focus on combination screens involving conventional chemotherapy agents with large non-specific small molecule libraries. Herein, we attempted to therapeutically leveraged the results of our chemogenomic data by designing a biology-driven drug combination screen. We first built a focused drug library with compounds chosen based on the following criteria: *i)* approved by the US FDA or in advanced clinical development, *ii)* considered potentially useful in treating brain tumors based on prior clinical and preclinical evidence, and *iii)* able to cross the blood-brain barrier. Then, using the biological and functional information obtained from our dual drug and siRNA screen, six out of the nine core vulnerabilities identified in GBM were targeted to perform a combination drug screen and quickly reveal potentially synergistic drug combinations. Our results indicated that synergism is rare with only 4.9% of the 528 tested pair-wise combinations described as potentially synergistic, in line with recent large-scale studies [38,39].

The MAPK pathway being one of the most commonly mutated oncogenic pathways in cancer, the therapeutic landscape of antitumour agents targeting this pathway has rapidly expanded, including several MEK inhibitors [40]. While studies have demonstrated the potential of MEK inhibitors in association with inhibitors of the PI3K/mTOR pathway [41,42], our drug combination screen with mTOR inhibitor Vistusertib indicated no potential synergy with the two tested MEK inhibitors Mirdametinib and Selumetinib. However, our results highlighted the potential synergistic interaction of CHK1 inhibitor Prexasertib with MEK inhibitors. Owing to its role in therapeutic resistance and oncogenesis, selective inhibitors of the cell cycle checkpoint kinase CHK1 are currently under pre-clinical and clinical investigations [43,44]. CHK1 has been shown to play a role in cell proliferation, radioresistance and Temozolomide-induced senescence in GBM [45,46]. Moreover, few studies showed that targeting the MEK pathway potentiates CHK1 inhibitor in multiple myeloma models, primary human glioma cells and a neuroblastoma cell line [47-49]. In line with these studies, using two GBM 3D spheroid models, we demonstrated the synergistic potential of combining Prexasertib with Mirdametinib, which warrants further investigation to confirm its therapeutic potential and clinical relevance in GBM.

Accumulating evidence demonstrate that aberrant epigenetic regulation of gene function plays a major role in carcinogenesis and tumor escape from therapy [50]. This has led to an increase in epigenetic drugs or epidrugs in development, including BET inhibitors (BETi) [51,52]. Owing to the interplay among various epigenetic processes, combining several epidrugs has emerged as a promising approach to anticancer therapy. Recent studies have provided mechanistic and pre-clinical evidence for the combination of HDAC inhibitors (HDACi) with BETi, even leading to the development of dual inhibitors [53-56]. Herein, our combination drug screen revealed that HDACi Panobinostat mostly antagonizes with all tested BETi in the murine GL261 cell line. While these unexpected results were confirmed in the GL261 spheroid model using Panobinostat and Birabresib, a synergistic effect was observed with the same drug combination in the U87 spheroid model. A recent study also indicated that HDACi combined with BETi was synergistic in two human GBM-sphere lines [57]. Our results thus suggest a cell-line dependance and warrant further molecular investigations to fully decipher the mechanism of action of the dual HDAC / BET inhibition.

Our results also highlighted the association of BET inhibition with AURKA targeting. The therapeutic potential of this combination has been previously shown in neuroblastoma models [58,59]. The synergistic effect has been notably linked to the inhibition of MYCN expression, as also demonstrated in an *in vitro* GBM study [60]. Moreover, by developing an integrative drug and disease signature approach, Stathias *et al*. recently identified JQ1 and Alisertib as a synergistic combination in GBM, which was validated in a subcutaneous model of GBM [61]. Nevertheless, as the high rate of attrition in cancer drug development is partly due to a lack of predictive preclinical models [62,63], findings need to be carefully evaluated using a range of model systems. Here, to confirm our results obtained with dual BET / AURKA inhibition on 3D spheroid GBM models and its therapeutic potential, we first developed an organotypic brain model in which murine GBM tumor progression was followed over time. These *ex vivo* tissue cultures are described as highly relevant models to study the evolution of pathologies and to test their response to different therapeutic strategies, including in brain diseases [64]. This co-culture system can keep brain tissues alive in incubation chambers mimicking natural conditions. Our results showed that the synergistic effect of Alisertib / Birabresib was maintained in this model and even exacerbated over time. To increase the complexity of our pre-clinical model and strengthen our results, we finally used a syngenic and orthotopic xenograft mouse model. We showed, for the first time, that the orally-administered drug combination significantly increased the median survival of GBM-bearing immunocompetent mice, without any observed side effects. As Alisertib has been already safely evaluated in a phase I for recurrent high-grade glioma patients [65], our preclinical data strongly support the therapeutic relevance of the Birabresib / Alisertib combination in GBM.

## CONCLUSION

Our findings highlight the prognostic value of RRM1 in GBM and the therapeutic potential of combining BET and AURKA inhibitors. Moreover, our data support the feasibility and efficiency of our approach to identify new disease-relevant targets by harnessing drug poly-pharmacology and using target deconvolution, and to therapeutically leverage these vulnerabilities by designing biology-guided drug combinations. This strategy could potentially be applied to any refractory cancers, thus opening major therapeutic avenues.

## ABBREVIATIONS

ATCC: American type culture collection;
AUC: area under the dose-response curve;
AURKA: aurora kinase A;
BET: bromodomain and extraterminal motif;
BETi: BET inhibitors;
CRISPR: Clustered Regularly Interspaced Short Palindromic Repeats;
DMSO: dimethyl sulfoxide;
FDA: federal drug administration;
GBM: glioblastoma;
GTeX: Genotype-Tissue Expression;
HDACi: Histone Deacetylase inhibitors;
IDH: isocitrate dehydrogenase;
MEK: mitogen-activated protein kinase;
MGMT: methyl guanine methyl transferase;
RRM1: Ribonucleoside-diphosphate reductase large subunit;
siRNA: small interfering Ribonucleic acid;
TCGA: cancer genome atlas;
TMA: Tissue microarrays;
TERT: telomerase reverse transcriptase;
WHO: world health organization.

## ACKNOWLEDGMENTS

We thank the ‘HiTS’ platform (Carine Derviaux; https://www.crcm-marseille.fr/equipes/plateformestechnologiques/hits-ipcdd/) for the help with the combination drug screening, and Elodie Berge for her technical assistance with the animal study. The authors would like to thank the AP-HM Tumor Bank (authorization number: AC2018-31053; CRB BB-0033-00097) for providing GBM tissue samples, and the ARTC-Sud patients association (*Association pour le Recherche sur les Tumeurs Cérébrales*). This work was supported by several research grants attributed to EP: Marie Sklodowska-Curie Fellowship from the European Research Council (FP7-PEOPLE-2013-IIF #626794), “Pépiniere d’Excellence” grant from the A*MIDEX Foundation of Aix Marseille University (funded by socio-economic partners), Translational Research grant from the French National Cancer Institute (PRT-K19-135, INCa), and research grants from Eva pour la Vie and Wonder Augustine and to MLG: post-doctoral fellowship from the ARC Foundation (#ARCPDF22020070002553). This work was also supported by the SIRIC Marseille (Integrated Oncology Research Site, INCa), the Canceropole PACA and several not-for-profit organisations (Association Léa, Association Cassandra, Le sourire de Lucie, Vocaliz’ et Nos p’tites Etoiles).

## CONTRIBUTIONS

EP and MLG designed the study, coordinated the work, analyzed the data and wrote the manuscript. ABJ performed functional validation of RRM1 and 2D and 3D cytotoxicity assays and the DepMap analysis, while MLG undertook the transcriptomic analyses of RRM1 and the validation experiments in 3D GBM spheroids with EP. RB performed *in vivo* experiments in orthotopic syngeneic model of GBM with the help of BM. MPM and PP realized the validation experiments in the organotypic model of GBM. GP designed the focused RNAi screen with EP, performed it and analyzed the results with EP and ABJ. TWF and GMA undertook the high-throughput drug screen and analysed the results with EP and ABJ while KM performed the combination drug screen with EP, MLG and XM. NBK, CC, OC, DB, NA, MC, ET and DFB were involved in study design and data analysis. DFB and CC supervised the protein expression analysis on patient samples. Finally, all authors read the manuscript and made comments to improve it.

