## Supplemental figures for "Combination drug screen targeting glioblastoma core vulnerabilities reveals pharmacological synergisms"

### SUPPLEMENTARY TABLES

*See excel file*

**Supplementary Table 1.** List of drug compounds for the primary screen.

**Supplementary Table 2.** Top 280 candidate compounds for the secondary screen.

**Supplementary Table 3.** List of 83 GBM killers.

**Supplementary Table 4.** List of interactors identified by target deconvolution.

**Supplementary Table 5.** List of 22 candidate genes.

**Supplementary Table 6.** Gene effect score.

**Supplementary Table 7.** Drug combination screen results.

### SUPPLEMENTARY FIGURES

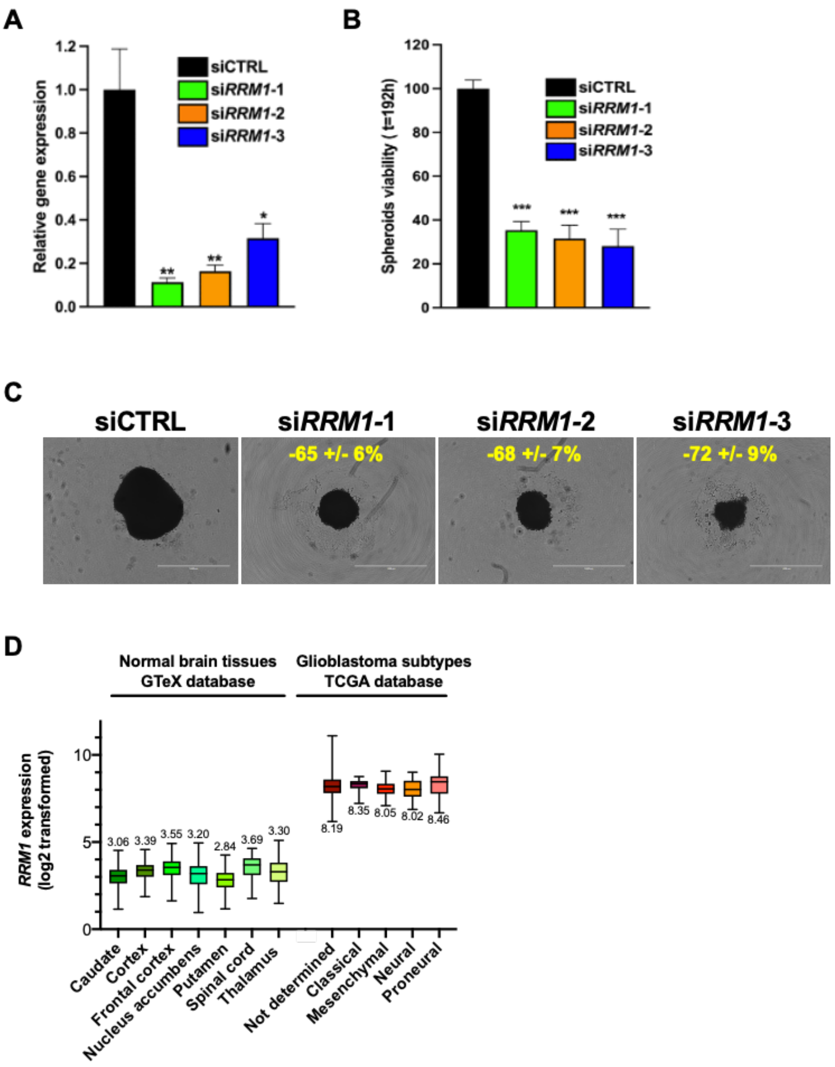


**Supplementary Figure 1.** (**A**) *RRM1* relative gene expression following 48h transfection of U87vIII cells with negative control siRNA and 3 different siRNA sequences targeting *RRM1*, as evaluated by qRT-PCR using *YWHAZ* as housekeeping gene. *Bars*, mean of at least 4 independent experiments; *Error bars*, S.E.M; *, $p<0.05$; **, $p<0.01$. (**B**) U87vIII tumor spheroid viability following siRNA transfection assessed by Alamar blue at the end point of the experiment (t=8 days). *Error bars*, S.E.M; ***, $p<0.001$. (**C**) Representative photographs of tumor spheroids at day 8 and percentages of decrease in spheroid growth compared to control (siCTRL) +/- S.D are included in insert. *Scale bar*, 1mm. (**D**) Relative *RRM1* gene expression in normal brain tissue (*green shades, left*) and GBM subtypes (*orange shades, right*), obtained from GTeX and TCGA databases, respectively (log2 transformed expression).

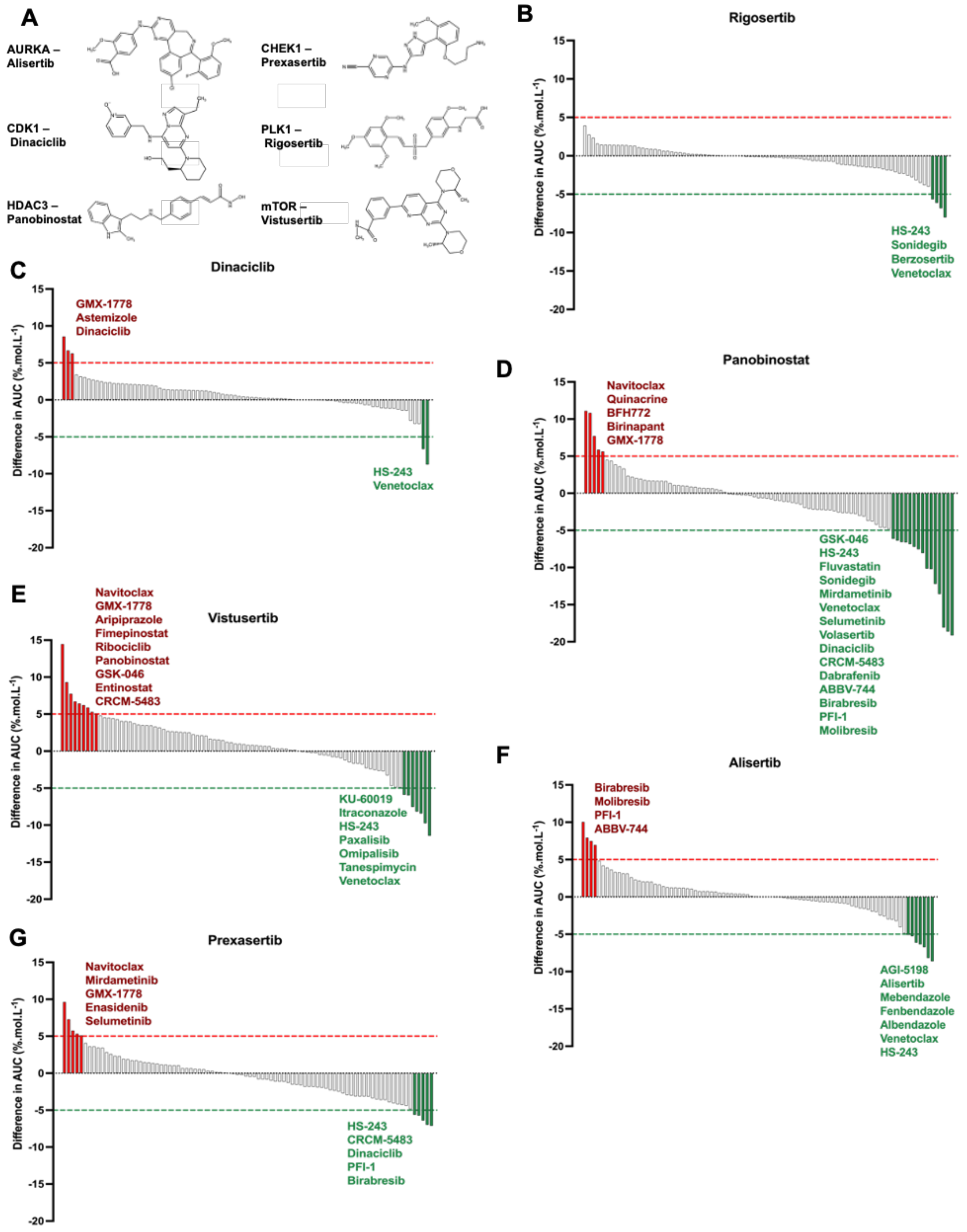


**Supplementary Figure 2.** (**A**) Representative molecule structures of AURKA inhibitor Alisertib, CDK1 inhibitor Dinaciclib, HDAC3 inhibitor Panobinostat, CHEK1 inhibitor Prexasertib, PLK1 inhibitor Rigosertib and mTOR inhibitor Vistusertib. (**B-G**) Fall plots show the AUC differences between the combinatorial treatment and the monotherapy condition obtained from our screen of a home-made drug library containing 88 drugs in GL261 cell lines. After 72h of drug incubation, cell viability was assessed using Cell Titer Glo®. Experiments were performed alone or in association with (**B**) Rigosertib 250nM, (**C**) Dinaciclib 5nM, (**D**) Panobinostat 10nM, (**E**) Vistusertib 250nM, (**F**) Alisertib 1µM and (**G**) Prexasertib 10nM.

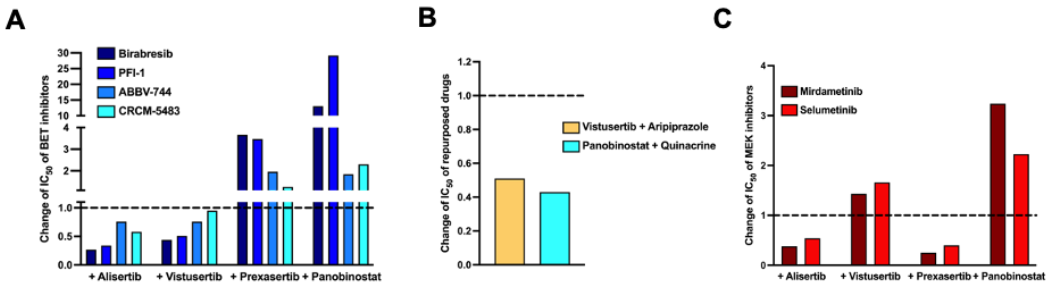


**Supplementary Figure 3.** Change in $IC_{50}$ values in GL261 cells when drugs are used in combination with Alisertib, Vistusertib, Prexasertib or Panobinostat, as compared with drug alone. Histogram representations for data obtained with (**A**) 4 epidrugs: Birabresib, PFI-1, ABBV-744 and CRCM5483, (**B**) 2 repurposed drugs Aripiprazole and Quinacrine in combination with Vistusertib and Panobinostat respectively, and (**C**) 2 MEK inhibitors: Mirdametinib and Selumetinib.

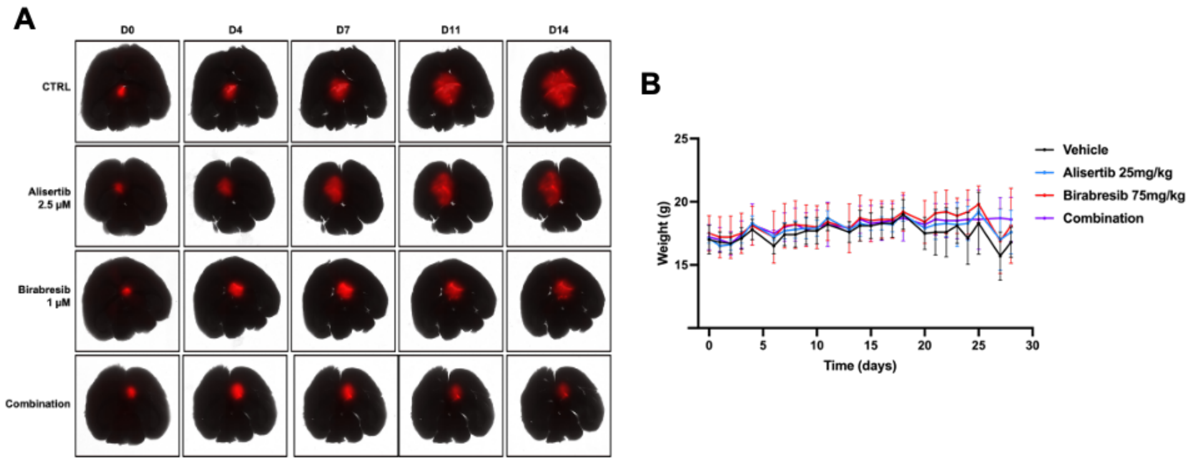


**Supplementary Figure 4.** (**A**) Representative pictures, acquired with the JuLi™ Stage live imaging system, of mtDsRed-expressing GL261 tumor micro-masses grafted in slices of healthy brain over time. *Scale bar*: 1mm. (**B**) Weight of murine GBM-bearing mice treated by oral gavage, one week after orthotopic injection of GL261 cells five time a week over three weeks, with vehicle only (1/10 DMSO in corn oil; *black*), Alisertib (75mg/kg; *blue*), Birabresib (25mg/kg; *red*) or the combination of both drugs (*purple*).
